# Single-step genome-wide association study for susceptibility to *Teratosphaeria nubilosa* and precocity of vegetative phase change in *Eucalyptus globulus*

**DOI:** 10.1101/2022.12.15.520574

**Authors:** Marianella Quezada, Facundo Giorello, Cecilia Da Silva, Ignacio Aguilar, Gustavo Balmelli

## Abstract

Mycosphaerella leaf disease (MLD) is one of the most prevalent foliar diseases of *E. globulus* plantations around the world. Since resistance management strategies have not been effective in commercial plantations, breeding to develop more resistant genotypes is the most promising strategy. Available genomic information can be used to detect genomic regions associated with resistance to MLD, which could significantly speed up the process of genetic improvement. In this study, we investigated the genetic basis of MLD resistance in a breeding population of *E. globulus* which was genotyped with the EUChip60K SNP array. Resistance to MLD was evaluated for resistance of the juvenile foliage, as defoliation and leaf spot severity, and for precocity of change to resistant adult foliage. Genome-wide association studies (GWAS) were carried out applying four Single-SNP models, a Genomic Best Linear Unbiased Prediction (GBLUP-GWAS) approach, and a Single-step genome-wide association study (ssGWAS). The Single-SNP and GBLUP-GWAS models detected 13 and 16 SNP-trait associations in chromosomes 2, 3 y 11; whereas the ssGWAS detected 66 SNP-trait associations in the same chromosomes, and additional significant SNP-trait associations in chromosomes 5 to 9 for the precocity of phase change (proportion of adult foliage). For this trait, the two main regions in chromosomes 3 and 11 were identified for the three approaches. The SNPs identified in these regions were positioned near the key miRNA genes, *miR156.5* and *miR157.4*, which have a main role in the regulation of the timing of vegetative change, and also in the response to environmental stresses in plants. Our results outlined that ssG-WAS was more powerful in detecting regions that affect resistance than conventional GWAS approaches. Additionally, suggest a polygenic genetic architecture for the heteroblastic transition in *E. globulus* and identified useful SNP markers for the development of marker-assisted selection strategies for resistance.

## 1 Introduction

*Eucalyptus globulus* Labill is widely planted in temperate countries mainly for the excellent quality of its wood for pulp production, which has a high market value in the paper industry^1^. However, the susceptibility of this species to several pests and diseases limits its use in commercial plantations^2^. Mycosphaerella foliar disease (MLD), caused by a complex of fungal species of the genus Mycosphaerella and Teratospaheria, is one of the most prevalent foliar diseases of *E. globulus* in natural forests and plantations worldwide^1^. *Teratosphaeria nubilosa* is considered one of the most virulent MLD species^3,4^. This pathogen infects predominantly juveniles and intermediate foliage and causes severe leaf spotting, premature defoliation and shoot blight^5^. Crown damage from MLD in young plantations can range from <10% to 80% which reduces the tree growth and survival, leading to concomitant loss in productivity of affected plantations^6,7^.

A number of silvicultural practices and management strategies have been proposed to minimize the effects of foliar diseases in Eucalyptus plantations. These include avoiding planting on high-risk endemic areas, the application of protectans and fungicides, as well using remedial fertilizer applications. However, these strategies have economic, environmental or operational constraints^2^. Thus, breeding for resistance to develop more resistant genotypes is the most promising strategy to effectively control MLD in commercial plantations^4,6,7^. Genetic variation on *E. globulus* for susceptibility to MLD has been identified between and within families, provenances and genetics groups^8,9^. Considering that this species is markedly heteroblastic, significant genetic variation in the timing of vegetative phase change has also been reported^6^. As young foliage is particularly susceptible to *T. nubilosa* infection, at least two mechanisms has been proposed to manage MDL disease: increase the resistance of juvenile foliage and increase the precocity change to the resistant adult foliage^7^. Several studies suggest that both mechanisms involved in foliar resistance are under moderate to strong genetic control in *E. globulus*^10^. But the underlying molecular mechanism and genetic architecture of MDL response has been relatively little investigated^11,12^.

Genome-wide association studies (GWAS) are a leading approach for complex trait dissection and identification of genomic segments or alleles that underlie phenotypic variation and thus can be used for breeding^13^. A common GWAS method in plants is the unified mixed linear model proposed by Yu et al.^14^, that accounts for relatedness at two levels: population structure and kinship. Since this method was computationally very intensive, a reparameterization of the MLM likelihood function was proposed^15^. Despite the statistical improvements, a typical feature of these GWAS methods is sequentially fitting each marker one at a time as fixed effects, resulting in a lack of power to map loci for quantitative traits^16^. This limitation has been overcome by methods that simultaneously fit high density genome-wide molecular markers by mixed linear models as GBLUP (Genomic Best Linear Unbiased Prediction). Following this strategy, marker’s effects are estimated by a linear transformation of genomic breeding values obtained from GBLUP^17^. One limitation of this method is that it only includes phenotypes of genotyped individuals. Generally in forest breeding populations, as in crops and livestock populations, only a fraction of individuals in a population are genotyped. Thus, the GBLUP methods were extended to include the information of non genotyped individuals in prediction analysis, which is the so-called Single-step genomic best linear unbiased prediction (ssGBLUP)^18^. The ssGBLUP approach is widely adopted for genomic prediction in livestock^19^ and recently applied in forest species^20,21^. The strength of this approach is the ability to jointly incorporate the information of all genotypes, observed phenotypes and pedigree information in one simple and single step. This procedure was extended to estimate the marker’s effects in an approach called Single-step GWAS (ssGWAS)^16^. Applying this method, marker effects and the p-values associated are estimated simultaneously for all markers while accounting for population structure using the pedigree and genomic relationship for all individuals in the population^22^.

The advances in next generation sequencing technologies allows the development of accessible high-throughput genotyping systems. Particularly for Eucalyptus species, in which marker number and density have been the limiting factors for a long time, dense sets of Single Nucleotide Polymorphism (SNP) genotyping platforms are now available^23^. The commercial Eucalyptus chip has been useful to understand the genetic basis of complex quantitative traits of interest, such as growth or wood properties in Eucalyptus species^24^. For example, in *E. grandis × E. urophylla* breeding populations, eight SNPs were described as main associations for growth traits, which were related to genes involved in cell wall biosynthesis^25^. Also, it has become possible to detect significant marker-trait associations for growth related traits in species poorly represented in the chip. Thus, a total of 87 SNPs were associated with growth and wood quality traits in *E. cladocalyx*, revealing associations with genes related to primary metabolism and biosynthesis of cell wall components^26^. Similarly, a study of *E. grandis × E. urophylla* hybrids detected 22 quantitative trait loci for growth and wood traits and four for *Puccinia psidii* rust disease resistance^27^. This resistance QTLs were positioned near the major QTLs detected in *E. grandis* for Myrtle rust (*Ppr1* locus). Recently, for the same fungal disease, 33 highly significant SNPs were detected in *E. obliqua*, identifying candidate defense response genes, one of them located within the *Ppr1* locus^28^. With the exception of Myrtle rust, GWAS analysis to investigate the genetic control and identify candidate defense genes for fungal diseases in Eucalyptus have not been implemented. For MLD resistance, a classical QTLs mapping using biparental crosses and 165 molecular markers detected two major QTLs that explained a large proportion of the phenotypic variance for *Mycosphaerella cryptic* severity^11^.

This work aimed to examine the genetic architecture of resistance to MLD caused by *T. nubilosa* in a breeding population of *E. globulus*. Although the Single-step GWAS strategy has received increasing attention in livestock species^19^, to the best of our knowledge, it has not yet been applied in forest tree species. Specifically, this study has two objectives: i) to compare the Single-SNP association GWAS, the GBLUP-GWAS and the Single-step GWAS strategies in a forest tree species, ii) to elucidate the genetic base of resistance and escape to *T. nubilosa* in *E. globulus*.

## 2 Material and Methods

### 2.1 Population and phenotypic data

This study was carried out using a progeny test of *E. globulus* established by Instituto Nacional de Investigación Agropecuaria (INIA) of Uruguay, in Lavalleja (Lat. 34 ° 110 S; Long. 54 ° 540 W; Alt. 206 m). The trial contain 194 open-pollinated families with a total of 4601 individuals. The experimental design was a randomized complete block design, with 3 replicates and eight-tree row plots.

When the trees were ~ 1 year old, there were several consecutive days of rain and high humidity, causing a severe infection of MLD^29^. Phenotypic responses to the natural infection were assessed at age 14 (coinciding with outbreak of MLD), 21 and 26 months. The response to disease was assessed using two parameters, the severity of leaf spots (SEV) and defoliation (DEF). Both were evaluated on the whole crown (juvenile and adult foliage) using a visual scale, recording the percentage of leaf area affected by spots (SEV) and the percentage of leaves prematurely abscised (DEF). The escape to disease was assessed as the precocity of vegetative phase change (ADFO), quantified as the proportion of the crown with adult foliage (petiolate, alternate, shiny green and pendulous leaves). The SEV was classified in percentages classes of 5%, whereas DEF and ADFO in intervals of 10%.

### 2.2 Genotyping

A total of 1008 trees were sampled for genotyping. DNA was extracted from leaf tissue of each tree using a standard CTAB protocol. Genotyping was performed using the Eucalyptus Illumina EUChip60K^23^ by GeneSeek (Lincoln, NE, USA). Of the 194 families in the trial, 179 have genotyped trees ranging from 4 to 8 trees per family. Individual genotypes were filtered by excluding samples with more than 10% of missing data across SNPs. The SNP data were then filtered by call rate > 90% and a minor allele frequency (MAF) > 0.01. The quality control of genotyping data was performed using qcf90 software^30^.

### 2.3 Population structure and linkage disequilibrium

The genetic structure of the breeding population was assessed by the Principal Components Analysis (PCA). Pairwise-estimates of Linkage disequilibrium (LD) were calculated as the squared correlation of allele counts for each pair of two SNPs (*r*^2^) within each of the 11 chromosome. The LD decay was estimated using the method described in Marroni et al.^31^. The pattern of LD decay was visualized plotting the *r*^2^ against the physical distance and the threshold of LD decay was identified at *r*^2^ =0.02. These calculations were performed using PREGSF90 of BLUPF90 family of programs^32^ and R software.

### 2.4 Statistical analyses

#### 2.4.1 Mixed model analysis

Linear mixed models were used to estimate the variance components of proportion of adult foliage, leaf spot severity and defoliation, considering the observed phenotypes and transformed into normal score within each block^33^. The linear mixed model was:

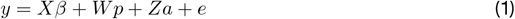

where *y* is the vector of phenotypes; *β* is the vector of fixed effects including the overall mean and block, with incidence matrix *X*; 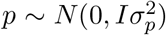 is the vector of random plot effects, with design matrix *W*; 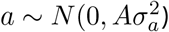 is the vector of random effects of individual trees (i.e., breeding values), with incidence matrix *Z*; and 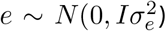 is the vector of residuals effects. The matrix *A* is the pedigree relationship matrix and 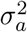 is the additive genetic variance. Genomic data were not included for variance components estimation.

Heritability of each trait was estimated as:

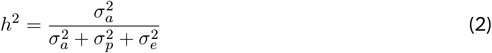

where 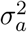 is the additive genetic variance, 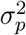 is the plot variance, and 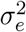 is the residual variance.

#### 2.4.2 Single-SNP models

The most common single-marker models were compared, with different combinations of population stratrification and genetic relatedness correction. First, a naive model, without any correction for population structure or relatedness was fitted independently for each SNP following the linear mixed model:

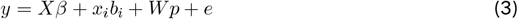

where *y* is the vector of phenotypes; *β* is the vector of fixed effects including the overall mean and block, with incidence matrix *X*; 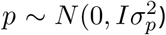 is the vector of random plot effects, with design matrix *W*; and 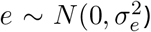 is the vector of residuals effects. The *x_i_* is a vector that contains the genotype for the *i^th^* SNP for each individual and *b_i_* is the *i^th^* SNP effect considered as fixed. A second model was fitted to account for population structure (model P), where the vector of fixed effects (*β*) of equation 3 include the first three principal components identified.

Alternatively, to account for family structure, a linear mixed model including a polygenic effect was tested (model K). This association model was fitted using the same naive model by adding a random effect utilizing a genomic relationship matrix as follow:

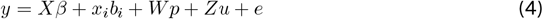

where *Z* is a desing matrix and *u* is the vector of polygene background effects (random), 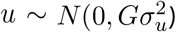. The rest of the components were previously defined (equation 3). The *G* matrix was estimated using the first method of VanRaden^34^:

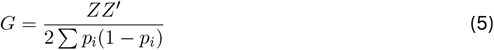

where *Z* is the matrix of gene content adjusted for observed allele frequencies and *p_i_* is the allele frequency of the *i^th^* SNP. To account for family and population structure simultaneously, a model K + P was fitted, where the vector of fixed effects (*β*) of equation 4 include the first three principal components. All Single-SNP models were implemented using the Sommer R package^35^.

#### 2.4.3 GBLUP-GWAS model

For the GBLUP-GWAS model, the follow mixed model was used:

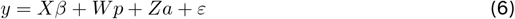

where every element is defined as before (equation 1), except for the vector of random effects of the individual trees (*a*) that follow a 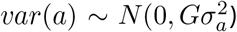. The matrix *G* is the genomic relationship matrix calculated using all the SNPs (equation 5).

Under this GBLUP-GWAS model, the vector of SNP effects was obtained from a linear transformation of the vector of breeding values in *a*. The vector of allele marker effects 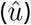 was calculated for all SNPs simultaneously following Gualdrón Duarte et al.^17^.

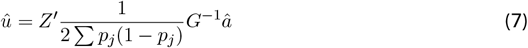

The variance of SNP effects and the p-values of each SNP effect were estimated following the formulas of Gualdrón Duarte et al.^17^.

#### 2.4.4 Single-step GBLUP association (ssGWAS)

The ssGWAS model is similar to the GBLUP-GWAS previously presented, but in this model all the phenotype and pedigree information of all assessed trees (genotyped and no-genotyped) was considered. Thus, the *G* matrix is replaced with the *H* matrix that combines both pedigree and genomic information.

The inverse of the relationship matrix that combines pedigree and genomic information (*H*^-1^) was derived by Aguilar et al.^18^ as:

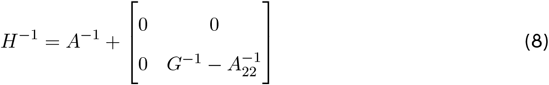

where *A*^-1^ and 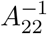 are the inverse of the pedigree relationship matrices for all individuals and only for the genotyped individuals, respectively, and *G*^-1^ is the inverse of the genomic relationship matrix.

The SNP effects were estimated as

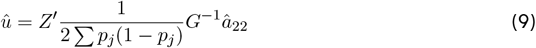

where 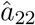 is a vector of genomic breeding values only for genotyped individuals. The variance explained for each marker and the p-values were obtained as suggested by Aguilar et al.^22^.

The GLUP-GWAS and ssGWAS models were fitted with POSTGF90 of BLUPF90 family software^36^. The GWAS results were plotted with CMplot R package^37^.

#### 2.4.5 Model comparison

Correction for multiple testing was applied to determine a threshold to indicated significant associations for all the six models implemented. At a genome-wide level, the Bonferroni correction at an alpha level of 0.05 was applied as a conservative criterion. In addition, a less stringent threshold for declaring significance was applied using the false discovery rate (FDR), with an alpha level of 0.05. Quantile-quantile (Q-Q) plots of observed and expected p-values were used to evaluate the control of population structure and genetic relationships in the different GWAS models. These plots were generated with qqman R package^38^.

### 2.5 Candidate genes and functional annotation

The physical position of the SNPs were defined according to the reference genome of *Eucalyptus grandis* (Phytozome Version 1.1 available on https://phytozome-next.jgi). Candidate genes were searched within 30 kb upstream and downstream of the significant SNPs. The size of this interval was chosen based on the typical LD present in the Eucalyptus populations and the SNPs density in our study (1 SNP per 28kb). The sequence of the candidate genes were extracted from the annotation file of *E. grandis* v1.1 .Particular miRNA related to heteroblasty, and their targets, were searched based on the annotation of Hudson et al.^12^ using blastn-short (BLASTN program optimized for sequences shorter than 50 bases). Functional annotation of genes associated with significant SNPs was done by protein homology search against the NCBI non-redundant protein sequences (NR) database^39^ using BLAST^40^. When possible, the best hit with functional annotation was used.

## 3 Results

### 3.1 Phenotypic variation

In this trial, the epidemic of *T. nubilosa* affected all the trees, which showed high variability for all the diseased-related traits. The mean and standard deviation for all the traits increased along with the increasing age of measurement, indicating a high prevalence of infection (Table 1). Though a wide range of variability was observed for the proportion of adult foliage, a high proportion of trees had only juvenile foliage, resulting in a distribution with an inflated proportion of zero counts in the three measurement ages. The disease damage traits (SEV and DEF) presented low but significant variability, following a normal distribution (Figure 1). Phenotypic correlations between measurement ages were positive and high for the proportion of adult foliage, and positive and moderate for defoliation. The disease damage traits, presented moderately positive correlation. The vegetative change to adult foliage (ADFO) was negatively correlated with response to diseased traits, confirming the mechanism of disease escape through early change to resistant adult foliage (Figure 1). The estimated pedigree-based heritabilities were moderate to high, varying from 0.33 for DEF_21 to 0.77 for ADFO_26 (Table 1).

**Figure 1:**
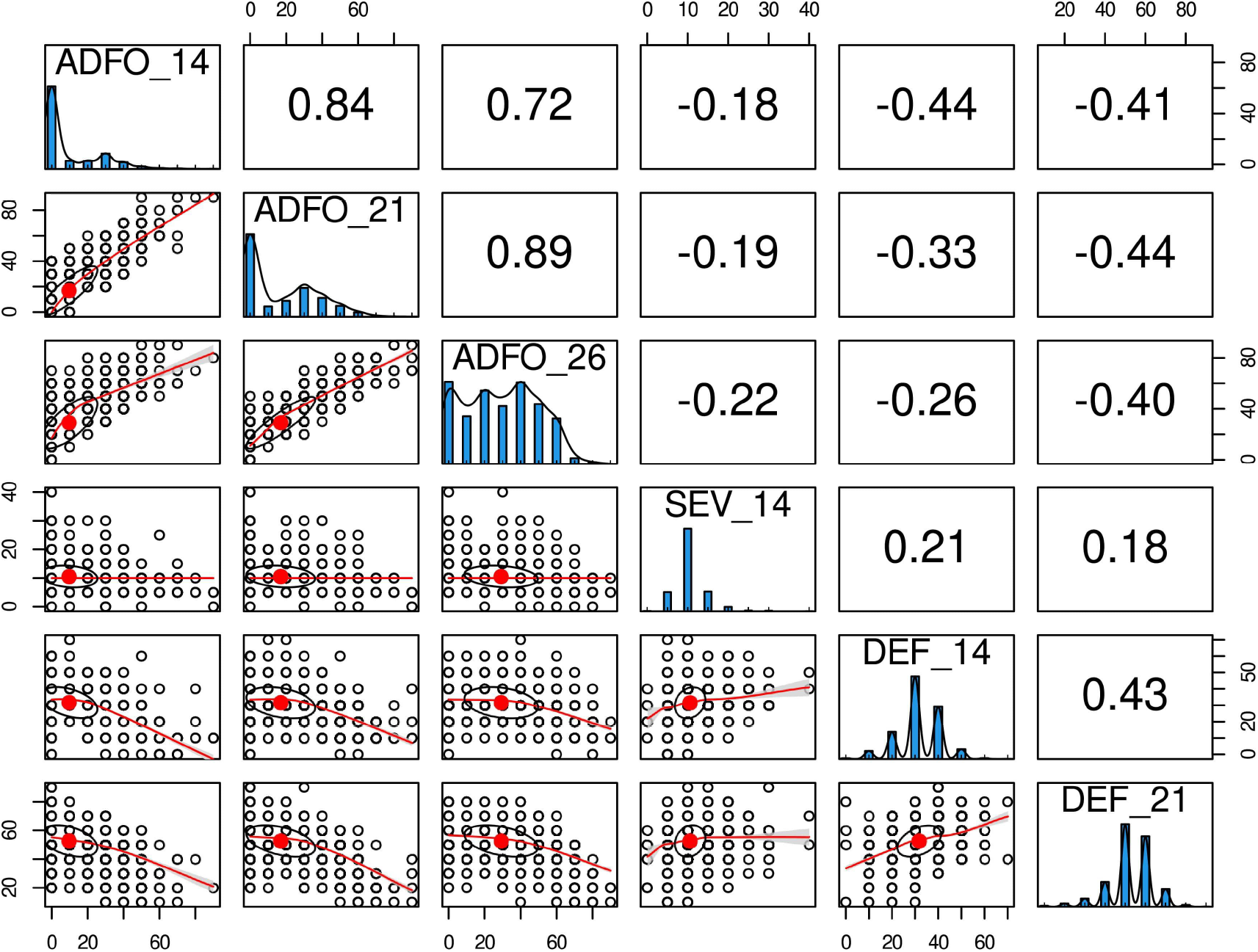
Phenotypic distribution and correlation of six diseased-related traits for *Eucalyptus globulus*. Histograms in the diagonal show the distribution of each trait. Scatter plots (lower off-diagonal) and Pearson’s correlation coefficient (upper off-diagonal) illustrate the underlying relationship between traits.

**Table 1:** Summary of assessed traits.

| Trait | Age of measurement<sup>a</sup> | Abbreviation | No. Obs | Mean (sd) | Range | $h^2$ |
| --- | --- | --- | --- | --- | --- | --- |
| Proportion of adult foliage | 14 | ADFO_14 | 3,853 | 9.76 (15.78) | 0-90 | 0.68 |
|  | 21 | ADFO_21 | 3,580 | 16.99 (19.18) | 0-90 | 0.66 |
|  | 26 | ADFO_26 | 3,447 | 29.14 (20.38) | 0-90 | 0.77 |
| Severity of leaf spots | 14 | SEV_14 | 3,711 | 10.53 (3.79) | 0-40 | 0.71 |
| Defoliation | 14 | DEF_14 | 3,711 | 31.62 (9.44) | 0-70 | 0.49 |
|  | 21 | DEF_21 | 3,581 | 52.73 (10.86) | 10-90 | 0.33 |

<sup>a</sup> in months

### 3.2 Population structure and linkage disequilibrium

A total of 19,992 markers and 961 trees were retained after quality filtering. The SNP markers were distributed throughout the 11 chromosomes of the *E. grandis* reference genome, with a range from 1,268 in chromosome 4 to 2,516 in chromosome 8 (Figure 2). The trend of Linkage Disequilibrium (LD) in the *E. globulus* population was analyzed across each chromosome. The average genowide *r*^2^ in this population was 0.032, with a LD decay at 19.1 kb using the significant threshold (*r*^2^ = 0.2) (Figure 3A). The LD decay varied across different chromosomes, with the most rapid LD decay observed for chromosome 5 (12.4 kb) compared with the slower rate on chromosome 9 (50.1 kb). The principal component analysis (PCA) showed the diversity and population structure present in the breeding population. The first two principal components explained 6.22% and 1.53% of the total variation, respectively. Based on these components, the majority of trees cluster into a major group, and different sub clusters can be identified (Figure 3B). Only the three first components that accounted for genetic variances greater than 1% were included in the association analysis to correct for population stratification.

**Figure 2:**
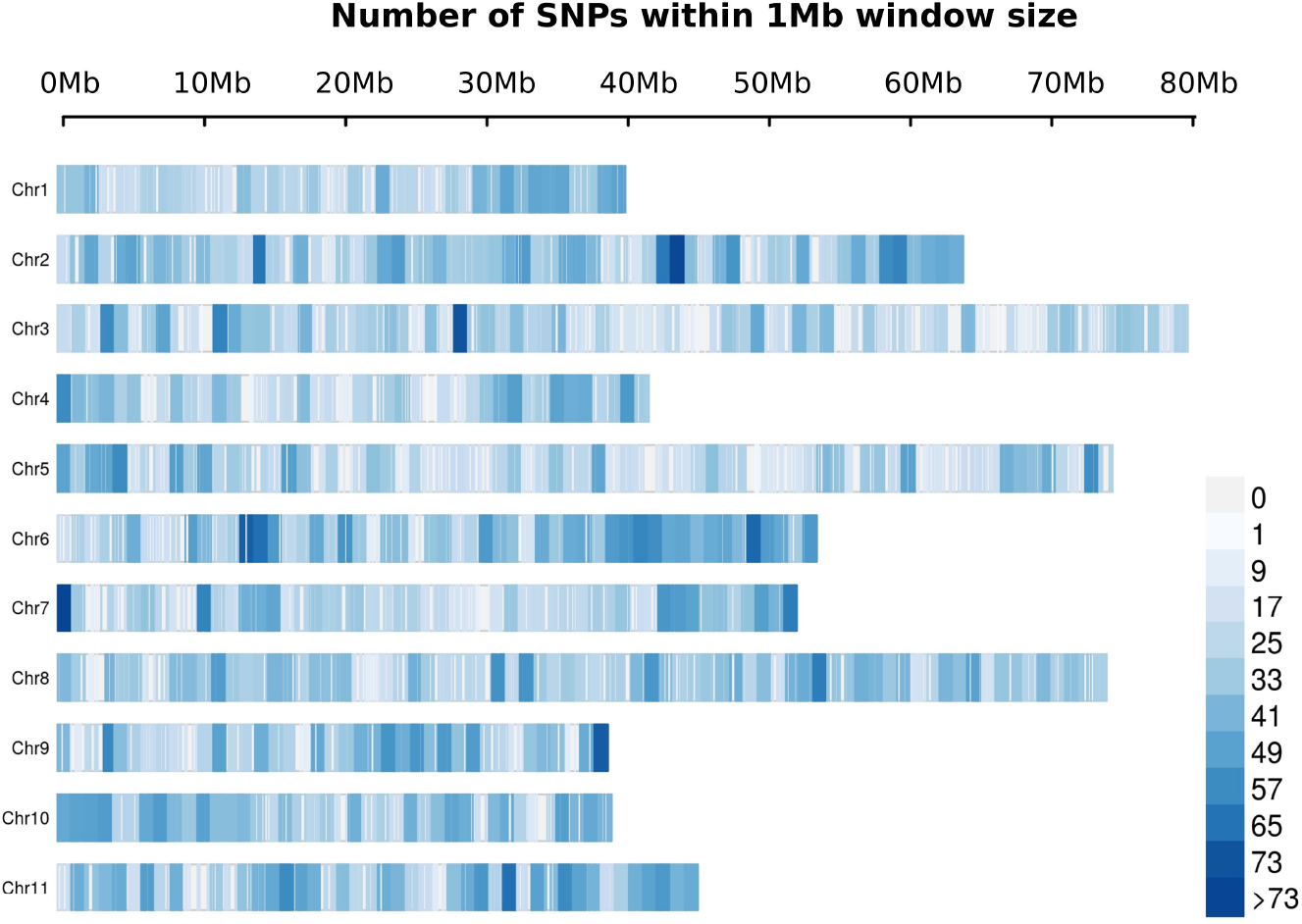
Distribution of filtered SNPs in a 1Mb windows across the *Eucalyptus grandis* genome. The x-axis represents the distance in Mb.

**Figure 3:**
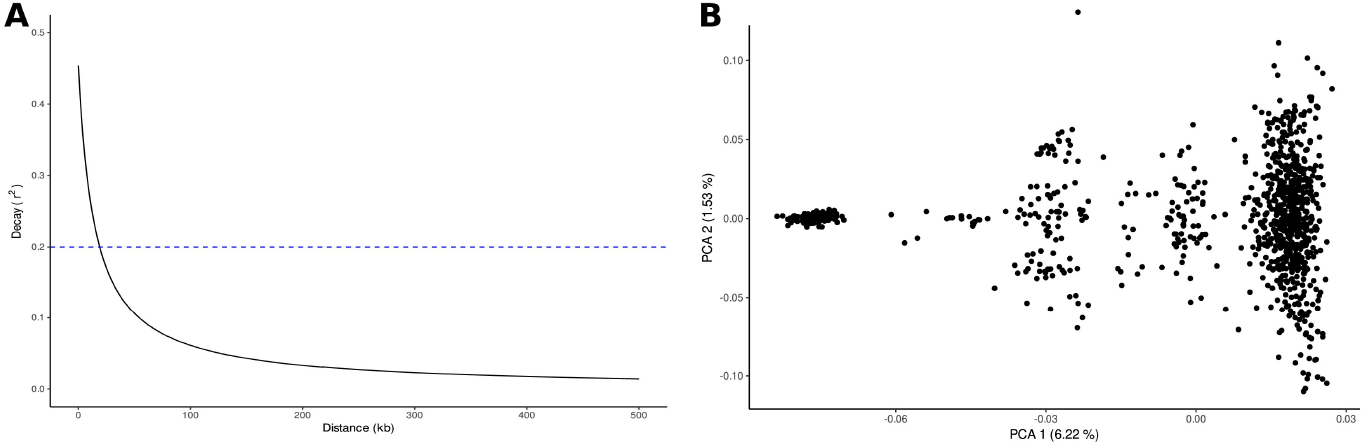
Population structure and linkage disequilibrium (LD) decay for the *Eucalyptus globulus* breeding population. (A) Linkage disequilibrium (LD) decay estimated by *r*^2^ (y-axis) plotted against physical distance in Mp (x-axis) along for a *E. globulus* population. Dashed line at *r*^2^ = 0.2 indicates the frequently used threshold of usable LD. (B) Principal Component analysis of 961 individual of the breeding population. PCA plot defined by first and second eigenvectors.

### 3.3 Association analysis

#### 3.3.1 Single-SNP models

The Single-SNP model without taking into account the population structure (naive model) resulted in the detection of a large number of associations for all traits (Table 2). Most of these were apparent false positives or spurious associations, by ignoring the multiple levels of relatedness between individuals. To control the population structure, the three first principal components were included in the model (model P). Although the number of associations slightly decreased, a high number of spurious signals were detected, due to family relatedness not being accounted for. The quantile-quantile plot (QQ-plot) obtained for the naive and model P, showed a great deviation for the observed and expected p-values, reflecting the inadequacy of these models (Supplementary Figure 1).

**Table 2:** Number of significant Single Nucleotide Polymorphism (SNP) associations for disease traits for all GWAS models evaluated. Significant associations using FDR (*alpha* = 0.05) correction. In parentheses number of significant associations using Bonferroni correction (*alpha* = 0.05).

| Trait | Single-SNP models |  |  |  | GBLUP-GWAS | ssGWAS |
| --- | --- | --- | --- | --- | --- | --- |
|  | naive | model P | model K | model K + P |  |  |
| ADFO_14 | 3,372 (260) | 3,282 (244) | 8 (3) | 6 (3) | 7 (5) | 26 (8) |
| ADFO_21 | 3,139 (193) | 3,081 (190) | 4 (2) | 4 (2) | 7 (2) | 25 (9) |
| ADFO_26 | 2,837 (165) | 2,797 (158) | 1 (1) | 1 (1) | 2 (1) | 15 (2) |
| SEV_14 | 104 (6) | 97 (4) | 0 (0) | 0 (0) | 0 (0) | 0 (0) |
| DEF_14 | 227 (2) | 222 (2) | 0 (0) | 0 (0) | 0 (0) | 0 (0) |
| DEF_21 | 372 (25) | 390 (29) | 0 (0) | 0 (0) | 0 (0) | 12 (1) |

When the random effects captured for the G matrix were included in the association models (models K and K+P), more p-values follow the uniform diagonal line in the QQ-plots (Supplementary Figure 1). Compared with the naive and model P, these models properly control false positive associations resulting from the within population family structure. As a result, including the G matrix resulted in a drastic reduction in the number of significant SNP associations. No significant associations were detected for SEV and DEF after correcting for multiple testing (Bonferroni and FDR at 5%) (Table 2). All the significant associations were detected for ADFO at the three measurement age. We found a total of 13 SNP-trait associations (8 unique SNPs) for the K model and 11 associations (6 unique SNPs) for the K + P model considering a FDR threshold of 5%, and 6 were also significant after the more stringent Bonferroni correction (5%), in both models. Moreover, both models did not show any difference, because all the SNPs detected in the K + P model were also significant in the K model. Thus, for this breeding population, the genetic matrix can account for genetic structure, with an negligible effect of the adjustment of principal components.

For ADFO, the significant SNPs were distributed in chromosomes 2, 3 and 11 (Supplementary Figure 2). The higher number of significant SNP associated were detected at the first age of measurement (14 months). It is not unexpected that the four and unique SNPs detected as significantly associated at the 21 and 26 months respectively, were also significant associated in the previous age measurement. Similar results were obtained using the observed variable and after the transformation to normal scores for all Single-SNP models tested, as well as for GBLUP-GWAS and ssGWAS models (results not shown).

#### 3.3.2 GBLUP-GWAS model

The GBLUP-GWAS model was performed for each variable to evaluate the equivalence with the Single-SNP model that accounts for genetic relationships in the G matrix (model K). For disease damage traits (SEV and DEF), no significant associations were declared using the multiple test correction (Bonferroni and FDR at 5%) (Table 2). Conversely, for ADFO the GBLUP-GWAS model identified 16 SNP-trait associations (8 unique SNPs) at the three measurement age at the most permissive level of FDR at 5%. Half of these associations were considered significant after Bonferroni correction (5%).

Of these 16 SNP-trait associations, 12 were previously identified by the Single-SNP model K and four were novel SNPs. The seven SNPs associated with the proportion of adult foliage at 14 months were also identified in the Single-SNP approach (model K) at the same measurement age (Figure 4). Similarly, all the SNPs associated at 21 and 26 months for the Single-SNP model K were detected in the GBLUP-GWAS model.

**Figure 4:**
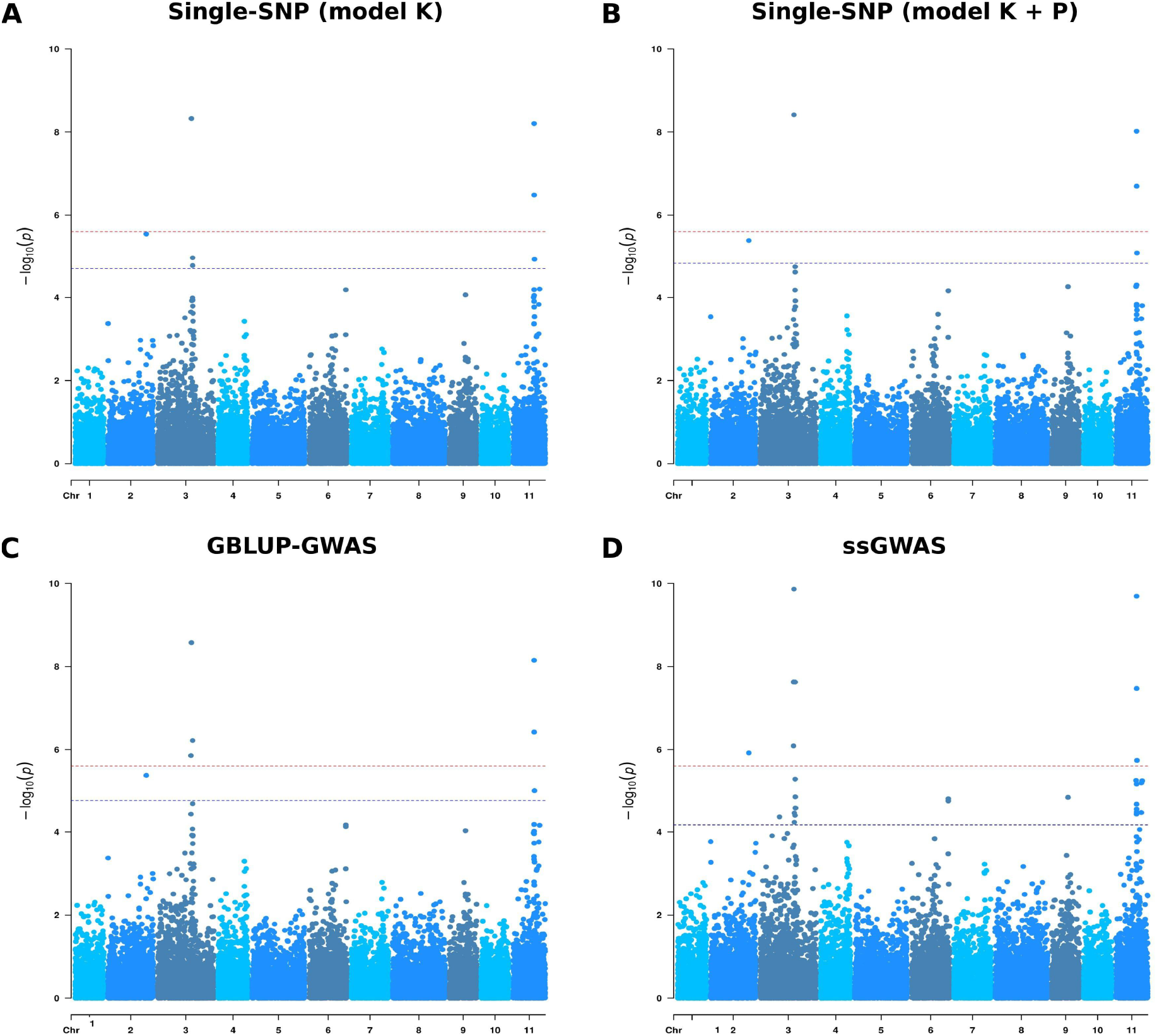
Manhattan plots for proportion of adult foliage measured at 14 months (ADFO_14) using three genome-wide association models. (**A**) Single-SNP model adjusted for kinship matrix (model K). (**B**) Single-SNP model adjusted for kinship matrix and population structure (model K + P). (**C**) GBLUP-GWAS model. (**D**) Single-step GBLUP association model. The x-axis represents SNP positions on the 11 *Eucalyptus grandis* chromosomes and the y-axis -log10 (p-values) from genotypic associations. The red horizontal line indicates the Bonferroni threshold (alpha = 0.05) and the blue horizontal line indicates a false discovery rate (FDR) at 5% threshold.

Associations for the proportion of adult foliage at 14 months were detected on chromosomes 2 (1 SNP), 3 (3 SNPs) and 11 (3 SNPs) (Supplementary Figure 3). The same regions in chromosomes 2, 3 and 11 were detected at 21 months, with six SNPs detected in both measurement ages. We found two SNPs, one in chromosome 3 and the other in chromosome 11, associated with ADFO at 26 months. These SNPs were associated with this escape to disease trait in the three measurement ages.

#### 3.3.3 ssGWAS model

The number of significant SNP markers detected in the Single-step GBLUP association approach for response and escape to disease-traits was larger than in the Single-SNP or GBLUP methods, which indicate a better performance of ssGWAS model (Table 2, Supplementary Table 1). Considering the FDR multiple test correction at 5%, we detected 66 SNP-trait associations and 38 unique SNPs for ADFO by the three measurement age. This approach identified 12 significant SNP associations for DEF at 21 months (Supplementary Table 1, Supplementary Figure 4). After the more conservative Bonferroni correction at 5%, 19 and one associations continue to be significant with ADFO and DEF traits, respectively. Once again, we found no significant association for SEV trait.

For ADFO, significant SNPs were associated across all chromosomes, except for chromosomes 1, 4 and 10. However, the most significant SNPs were identified in chromosomes 3 and 11 in the three measurement ages, indicating the presence of few genomic regions controlling the precocity of adult foliage change (Figure 5).

**Figure 5:**
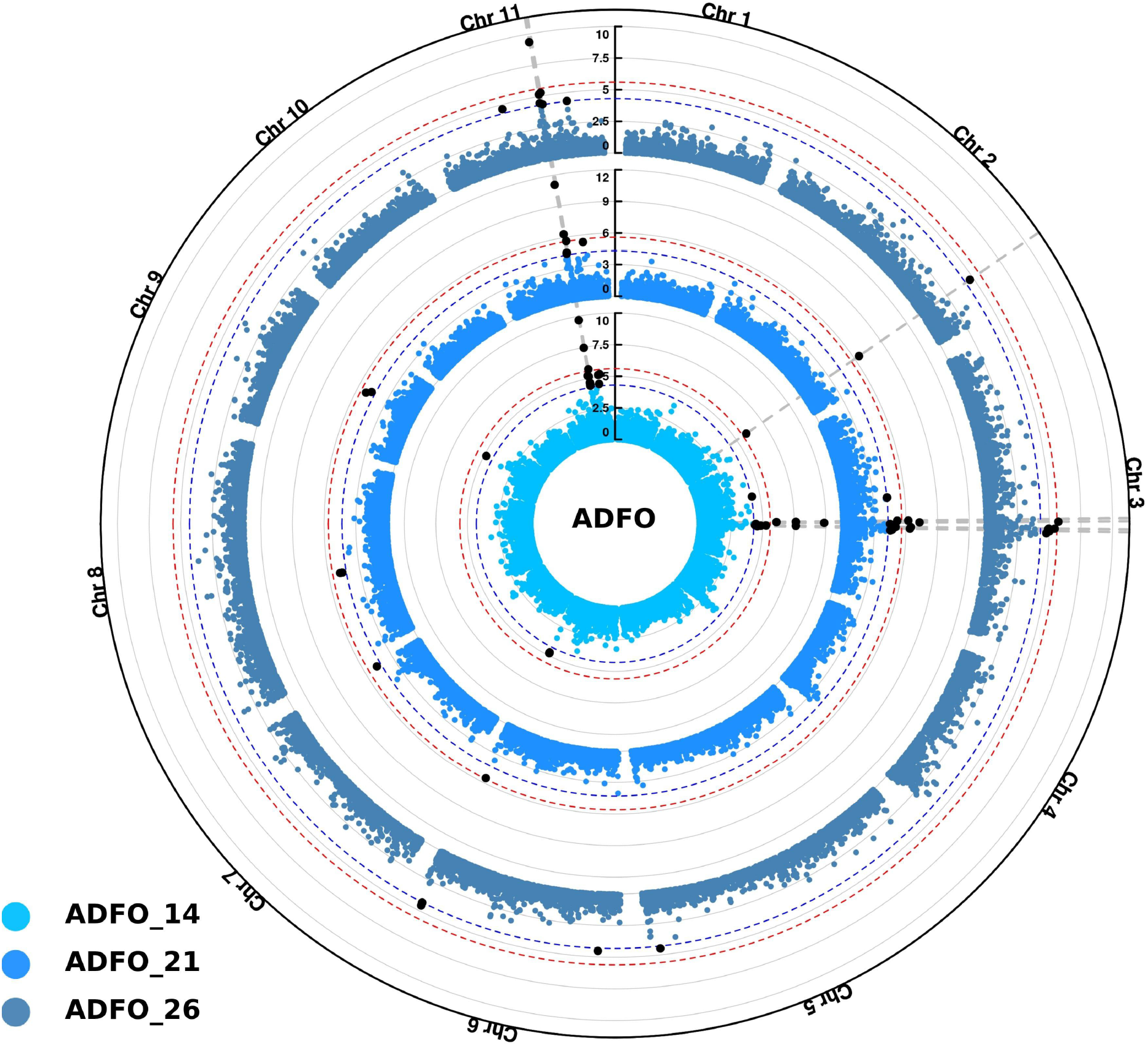
Circular manhattan plots for proportion of adult foliage (ADFO) using Single-step GBLUP association model (ssGWAS). Inner, middle and outer layers represent the measurement ages at 14, 21 and 26 months, respectively. The Bonferroni (alpha = 0.05) and false discovery rate (FDR) threshold at 5% are represented by the red and blue circle in each layer, respectively. Vertical dashed lines highlight common SNPs identified among the three measurement ages. Significant SNPs identified after multiple test corrections are in black.

The first peak identified on chromosome 3 spans a region of ~ 2 Mb, defined in accordance with SNPs overlapping at the three age measurement. The high number of significant SNPs was associated with the first age of measurement (14 months), namely nine SNPs located in this region, the most significant with a -log10(p-value) of 9.86 (position 47 949 673). Inside this area, the six SNPs significantly associated at 21 months, were also detected at the measurement age of 14 months. Additionally, the five SNPs found at the 26 months, were also detected in the two previous measurement ages. The SNPs detected in this region explained the 1.14, 1.64 and 0.44 of the total genetic variance for each age at measurement of 14, 21 and 26 months, respectively.

A small region of ~ 200 kb was also identified in chromosome 11 significantly associated with this trait. This region encompasses 7, 5 and 3 SNP for measurement at 14, 21 and 26 months. Once again, the high number of SNPs was identified at the first measurement age (14 months), and all the markers identified at 21 and 26 months were also detected in the previous measurement age. In this region, the SNP in the position 29 564 447 was the most significant, with -log10(p-values) values of 9.69, 11.07 and 9.36 for 14, 21 and 26 months, respectively. The SNPs inside this region represented 1.14, 0.94 and 0.73 of the genetic total variance at the three measurement age, respectively.

Additionally, other significant signals were identified in chromosome 2, with one SNP (position 54 236 535) detected at the conservative threshold of Bonferroni correction at 14 and 21 months, and with the FDR threshold at 26 months. For chromosome 6, two SNPs located 100 kb apart were identified with the less stringent FDR correction for the three measurement ages (Figure 5).

The ssGWAS model was able to detect 12 SNPs in chromosomes 3, 6, 8 and 11 associated with the defoliation measured at 21 months. However, no significant associations were detected for the previous measurement age (14 months). As expected by the correlation between defoliation and the proportion of adult foliage traits, six common SNPs between these traits at the same age (21 months) were observed. Thus, two SNPs identified in chromosome 3 and four in chromosome 11, correspond to the significant regions detected for ADFO.

### 3.4 Candidate Genes for precocity of vegetative phase change

Of the 38 unique significant SNPs identified for heteroblasty, two of them did not have any associated gen. A total of 133 genes were associated with the rest of the significant SNPs, of which 131 were protein coding. Two key miRNA genes, *miR156.5* and *miR157.4*, were found in chromosome 3 and 11, respectively (Table 3). The significant SNPs associated with these genes were at 21 kb and 25 kb for *miR156.5* and *miR157.4*, respectively. None of the *miR156* and *miR157* target proteins were found.

**Table 3:**
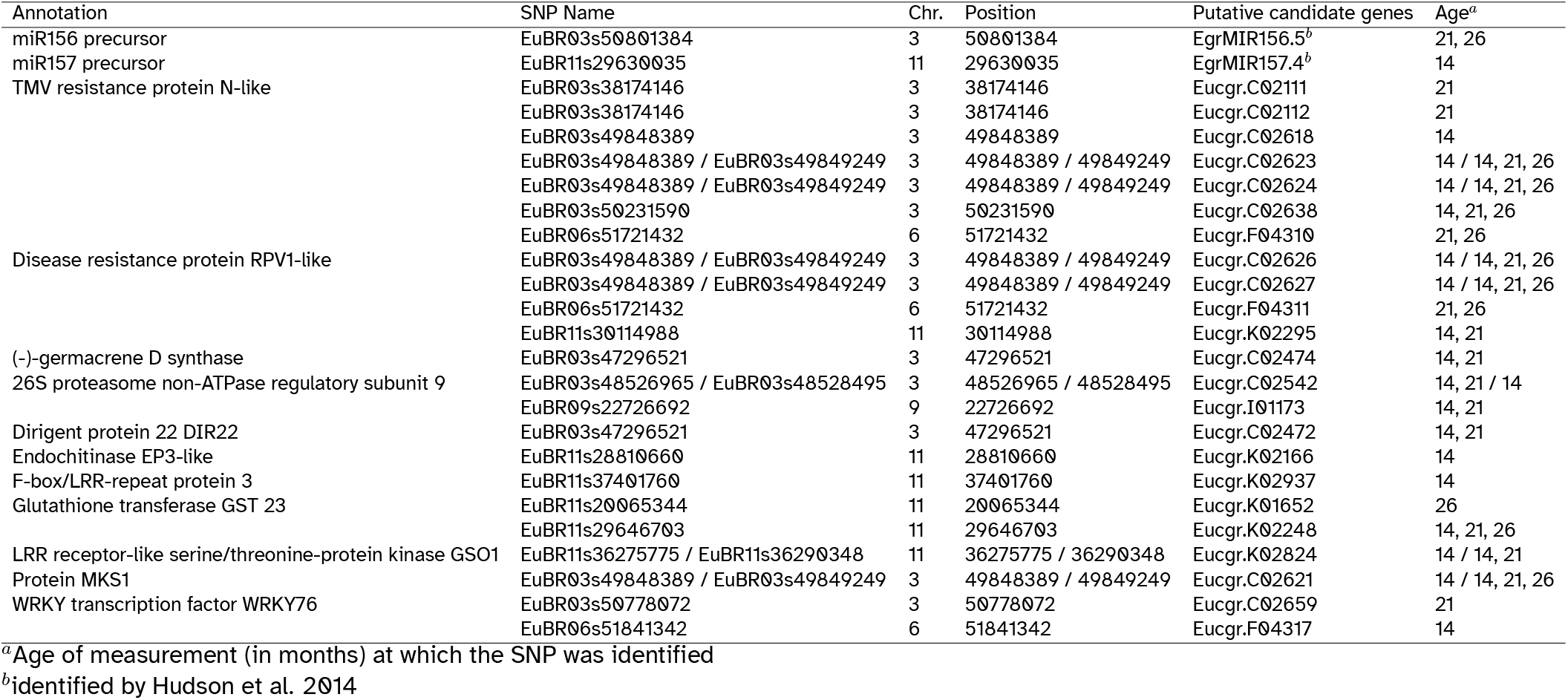
Details of genes related with biotic stress resistance linked to the most significant Single Nucleotide Polymorphism (SNP) associated with precocity of vegetative change (proportion of adult foliage - ADFO) in *Eucalyptus globulus* identified with the ssGWAS model

Out of the 131 protein coding genes, 20 could not be assigned a specific functional annotation. In total 95 different functions were found (Supplementary Table 2). A total of 11 proteins related to resistance to biotic stress encoded by 22 different genes were the most interesting ones: 26S proteasome non-ATPase regulatory subunit 9 (Eucgr.C02542), dirigent protein 22 (Eucgr.C02472), disease resistance protein RPV1-like (Eucgr.C02626, Eucgr.C02627, Eucgr.F04311, Eucgr.K02295), endochitinase EP3-like (Eucgr.K02166), F-box/LRR-repeat protein 3 (Eucgr.K02937), (-)-germacrene D synthase (Eucgr.C02474), glutathione transferase GST 23 (Eucgr.K01652, Eucgr.K02248), LRR receptor-like serine/threonine-protein kinase GSO1 (Eucgr.K02824), MAP kinase 4 substrate 1 - MKS1 (Eucgr.C02621), TMV resistance protein N-like (Eucgr.C02111, Eucgr.C02112, Eucgr.C02623, Eucgr.C02624, Eucgr.C02638, Eucgr.C02618, Eucgr.F04310), WRKY transcription factor WRKY76 (Eucgr.C02659, Eucgr.F04317) (Table 3).

## 4 Discussion

In a GWA study, for any population of a given genetic background, the ability to detect associations is a function of several parameters that include the phenotypic variance, the heritability and the complexity of the trait. In the current study, we show significant phenotypic variance for resistance to MLD for all the traits assessed in a breeding population of *E. globulus*. The heritabilities were high (*h*^2^ > 0.65) for ADFO and SEV, whereas moderate values were observed for DEF. These estimates were similar or higher to those reported in previous studies carried out for equivalent resistance or damage traits of *Teratosphaeria* fungal diseases in *E. globulus*. For example, heritabilities for severity to *T. nubilosa* and *T. cryptica* ranging from 0.12 to 0.60^7,9,41–43^ whereas values between 0.43 to 0.74 have been reported for proportion of adult foliage^6^. Regarding the genetic complexity of the MLD resistant traits in *E. globulus*, two major genomic regions which explained a large proportion of the phenotypic variance have been previously identified, and an oligogenic or major gene control proposed for these traits^11^. However, recent studies show that resistance mechanisms for other diseases in Eucalyptus involve multiple interacting loci of variable effect, according to a quasi-infinitesimal model^27,44^. Specifically for heteroblastic transition, although major genes have been identified in various species, still have a rather limited understanding of the number of loci involved in the regulatory pathway of this process^12^. Thus, for the disease-related traits analyzed here a quantitative nature is expected, which means that the phenotypic variation observed can be dependent on more than one major QTLs and many significant marker-trait associations would be identified.

In a first approximation to the genetic architecture to MLD resistance, we applied the widespread mixed model approach proposed by Yu et al.^14^. We used the PCA scores to control for population stratification and the genomic relationship matrix (G) to control for close family relatedness. As expected, when the background genetic structure is not properly accounted for in the model, a high rate of false positive marker-trait associations are observed (Supplementary Figure 1). Our results, in line with previous association studies in Eucalyptus, demonstrated that the inclusion of family relatedness correction in association models drastically reduces the number of false-positive associations^25,45^.Thus, for breeding populations consisting of families with cryptic relatedness, such as open-pollinated populations, the family structure (G matrix) efficiently captures the variations caused by genetic structure. The adjustment of using the PCA scores may be not adequate, and would be only necessary in populations with high levels of subrace differentiation or with different geographic origins^46^.

In the standard GWAS approach, each marker is tested independently for an association to the trait, included as a fixed effect in the model. For this method, two main drawbacks have been identified. First, the magnitude of the effects by the marker-trait associations identified are largely overestimated, caused mainly by the limited sample sizes and/or the use of the same data set for discovery and parameter estimation^47^. Second, the effect of one marker is estimated ignoring the rest of the genome, causing confusion about the number and location and also contributing to the upward bias of estimated effects of loci identified^48^. The alternative approach that fits all markers simultaneously as a random effect, GBLUP-GWAS, is becoming popular due to having much in common with genomic selection^34^. In our study, the GBLUP-GWAS and the Single-SNP approach (model K) show similar results in the marker-trait associations identified, where 75% of the markers identified in the GBLUP model were previously identified in the Single-SNP model. When the SNPs effects for this common SNPs were compared, our results agree with previous results that show an upwardly biased estimation under a fixed single-marker model^48^.

For GWA studies, the use of more phenotypic and genotypic data improves power and resolution^13^. Thus, the ssGWAS method that include the phenotypes of all genotyped and non-genotyped individuals as well the pedigree information, have been proven to provide more consistent solutions and increasing accuracy than the standard GWAS approach^16^. Additionally, as demonstrated by Mancin et al.^49^ using simulated data, ssGWAS can efficiently account for population structure, like GWAS-GBLUP models. Although this approach was successfully applied in GWA studies in domestic animals^19^, to our knowledge, this is the first report of applying ssGWAS in a forest tree species. In this study, the ssGWAS approach identified 66 and 12 significant SNP-trait associations for the ADFO and DEF traits, respectively. Although, our results show a general agreement between the Single-SNP and GBLUP, the ssGWAS demonstrated superior performance by detecting new significant SNPs in chromosomes that had not been identified in the Single-SNP or GBLUP approaches. In addition, although the same genomic regions were identified with all models, the ssGWAS models resulted in clear peaks with higher -log10(p-values). Therefore, the ssGWAS appears to be a promising method to the dissection of complex traits in populations when only a fraction of the population is genotyped, taking advantage of the available phenotypic information of individuals non-genotyped.

Most studies addressing disease resistance in forest trees have mainly assessed the disease’s severity as the presence of leaf spots and/or the necrotic lesions present on the leaves. The high heritability found for severity of leaf spots (SEV) in agreement with previous reports, broadens the probability of detecting a gene of large effect. However, large heritability does not imply a direct relationship with the number, effect or proportion of the genetic variance of the genomic regions identified^50^. Our outcome can be explained by the low phenotypic variability of this trait in this population. Freeman et al.^11^ showed that the number of resistance QTLs detected in three breeding populations of *E. globulus* was strongly correlated with the variability within each population. In this study we also evaluated the degree of defoliation (DEF) as a descriptor of the impact of the MLD infection. For this trait, assessed at 21 months of age, we found 12 significant associations, although only one was significant after Bonferroni correction (Table 1). The low number of significant associations can be partially explained by the moderately-low heritability of this trait, reflecting the superior performance of the ssGWAS approach relative to the standard GWAS approaches.

The vegetative phase change in plants involves many morphological changes, well differentiated in many tree species as *E. globulus*. The mechanism to the juvenile to adult transition is under a strong genetic control with the same regulatory mechanisms in both annual herbaceous plants and perennial trees^51^. Although the understanding of the genetic control of this transition remains limited, it is mediated by environmental and endogenous signals^52^. In *E. globulus*, precocious vegetative change was evidenced in response to abiotic stress (drought, salt and high winds) in a natural coastal population^53^. Also, it is proposed as a mechanism involved in disease resistance, anticipating the timing of vegetative change to the resistant adult foliage. For precocity of vegetative change, measured as the proportion of adult foliage in this study, the ssGWAS model was able to identify the highest number of significant trait-SNP associations in this breeding population. For the three measurement ages, the same SNPs were identified as the markers that explained the highest genetic variation, which allowed identify two main significant regions in chromosome 3 and 11. Within those regions, we identified the *miR156.5* and *miR157.4* respectively, both of which target DNA-binding transcription factors that regulates the SQUAMOSA PROMOTER BINDING PROTEIN-LIKE (SPL) gene family. The *miR156* has been recognized as a master regulator of vegetative change in plants, promoting juvenility due to high expression in seedlings that decrease during development^52,54^. Our results support similar reports from bi-parental QTLs mapping studies that identified *miR156.5* as the candidate gene causing difference in the timing of phase change between precocious and normal natural populations of *E. globulus*^12^. Furthermore, emerging studies suggest that *miR156* is probably an integral component of the *miRNA* response to all environmental stresses in plants^55,56^. Specifically for fungal pathogen defense response, *miR156* targets the plant disease resistance proteins (NBS-LRR proteins) in *Populus trichocarpa* infected with stem canker pathogen^56^. Additionally, precursors of *miR156* have also been reported located near the main QTL identified for *Puccinia* rust resistance (*Ppr1)*, suggesting a main role in the genetic control of plant resistance^44^.

We also identified various genes proposed to be involved in fungal pathogen resistance through different mechanisms. For example, the 26S proteasome non-ATPase regulatory subunit 9 is part of the ubiquitin/26S proteasome system, known to be involved in almost every step of the defense mechanisms in plants, regardless of the type of pathogen, regulating key cellular processes through protein degradation^57^. The dirigent protein 22 (DIR22) is part of a gene family involved in plant defense by their role in the dynamic reorganization of the cell wall through the formation of lignans and production of defense-related compounds^58^. Numerous studies have shown that the expression levels of DIRs genes are enhanced during the late stages of fungal infection (e.g., *Colletotrichum gleosporioides, Botrytis cinerea, Fusarium oxysporum*) in various plant species, increasing antifungal responses^59–61^. Glutathione transferase GST23 belong to a super-family of multifunctional proteins in plants, being one of their most important roles the detoxification of fungal toxins in the response against biotic stresses^62^. The expression of GSTs is markedly induced during fungal infection, leading to enhanced resistance to the pathogen^63^. The (-)-germacrene D synthase is an enzyme involved in the volatile terpenoid biosynthesis. It has been reported that the downregulation of these genes alters monoterpene levels leading to resistance against biotic stresses at a global level, including defense responses to fungus infection^64^. Lastly, the endochitinase EP3-like genes encode a protein involved in catalyzing the hydrolytic cleavage in chitin, the prominent wall component of fungus, and are induced by pathogen attack and other biotic stresses^65^.

Finally, we identified genes involved in pathogen defenses’ regulation and signaling mechanisms. MAP kinase 4 substrate 1 (MKS1) has been reported to be pivotal in signaling basal defense responses^66^, whereas the WRKY transcription factor WRKY76 is involved in the regulation of the plant defense signaling pathway^67^. In addition, many plant disease resistance proteins (NBS-LRR proteins), which trigger signal transduction leading to the induction of defense responses^68^, were found. For instance, the disease resistance protein RPV1 is known to confer resistance to oomycete and fungal pathogens in cultivated grapevines (*Vitis vinifera)^69^*. LRR receptor-like serine/threonine-protein kinase GSO1 was reported to be related to the resistance of *Nicotiana tabacum* to the oomycete *Phytophthora parasitica*^70^. TMV resistance protein N-like was reported to be involved in the resistance in potato to the fungus *Synchytrium endobioticum*^71^, and in peanut to two serious fungal foliar diseases^72^. F-Box/LRR repeat protein was reported to be activated by fungal pathogens (*Cladosporium fulvum, Puccinia striiformis*) in different plant species in order to regulate defense responses^73^.

### 4.1 Conclusions

In the present study, we carried out GWAS for traits related to response and escape to diseases in a breeding population of *E. globulus* only partially genotyped, and assessed the advantage of including phenotypes from relatives without genotypes in a single-step procedure. We compare several GWAS models that differ in the correction for population structure, the statistical approach to fit the markers and the impact of including combined pedigree, genomic and phenotypic information. Several SNP-trait associations were detected for the precocity of vegetative phase change (proportion of adult foliage). The same SNPs were identified at different measurement ages for the different strategies, providing validation of the results for these specific loci. Additionally, our results suggest that ssGWAS is a powerful model in detecting regions associated to the escape to MLD, showing a greater number of significant SNP associations than Single-SNP and GBLUP-GWAS models. Associations were detected near the miRNAs that regulate the timing of vegetative phase change, consistent with results from other studies about heteroblastic transition in trees and annual species. Thus, our research supports the hypothesis that this is a complex trait determined by a large number of genes with diverse biological functions, that can also include a mechanism of response to biotic stress, such as fungal resistance. Although the identified putative candidate genes will require future validations, they provide a better understanding of the genetic architecture of vegetative change in Eucalyptus.

## Data Availability Statement

The phenotypic and genotypic data analyzed in this study are not publicly available. The data are available upon reasonable request to the corresponding author and with permission of INIA.

## Author Contributions

MQ, IA, and GB contributed to conception and design of the study. GB designed the field experiment, collected the phenotypic data and provided the marker data. MQ performed the statistical analysis under the supervision of IA. FG and CS performed the candidate gene analysis and wrote this section of the manuscript. MQ wrote the final version of the manuscript. All authors contributed to manuscript revision, read, and approved the submitted version.

